# *Mycobacterium tuberculosis* requires the outer membrane lipid phthiocerol dimycocerosate for starvation-induced antibiotic tolerance

**DOI:** 10.1101/2022.07.07.499240

**Authors:** Alisha M. Block, Sarah B. Namugenyi, Nagendra P. Palani, Alyssa M. Brokaw, Leanne Zhang, Kenneth B. Beckman, Anna D. Tischler

## Abstract

Tolerance of *Mycobacterium tuberculosis* to antibiotics contributes to the long duration of tuberculosis (TB) treatment and the emergence of drug-resistant strains. *M. tuberculosis* drug tolerance is induced by nutrient restriction, but the genetic determinants that promote antibiotic tolerance triggered by nutrient limitation have not been comprehensively identified. Here, we show that *M. tuberculosis* requires production of the outer membrane lipid phthiocerol dimycocerosate (PDIM) to tolerate antibiotics under nutrient-limited conditions. We developed an arrayed transposon (Tn) mutant library in *M. tuberculosis* Erdman and used orthogonal pooling and transposon sequencing (Tn-seq) to map the locations of individual mutants in the library. We screened a subset of the library (~1,000 mutants) by Tn-seq and identified 32 and 102 Tn mutants with altered tolerance to antibiotics in stationary phase and phosphate-starved conditions, respectively. Two mutants recovered from the arrayed library, *ppgK*::Tn and *clpS*::Tn, showed increased susceptibility to two different drug combinations in both nutrient-limited conditions, but their phenotypes were not complemented by the Tn-disrupted gene. Whole genome sequencing revealed single nucleotide polymorphisms in both the *ppgK*::Tn and *clpS*::Tn mutants that prevented PDIM production. Complementation of the *clpS*::Tn *ppsD* Q291* mutant with *ppsD* restored PDIM production and antibiotic tolerance, demonstrating that loss of PDIM sensitized *M. tuberculosis* to antibiotics. Our data suggest that drugs targeting production of PDIM, a critical *M. tuberculosis* virulence determinant, have the potential to enhance the efficacy of existing antibiotics, thereby shortening TB treatment and limiting development of drug resistance.

**IMPORTANCE:** *Mycobacterium tuberculosis* causes 10 million cases of active TB disease and over 1 million deaths worldwide each year. TB treatment is complex, requiring at least 6 months of therapy with a combination of antibiotics. One factor that contributes to the length of TB treatment is *M. tuberculosis* phenotypic antibiotic tolerance, which allows the bacteria to survive prolonged drug exposure even in the absence of genetic mutations causing drug resistance. Here we report a genetic screen to identify *M. tuberculosis* genes that promote drug tolerance during nutrient starvation. Our study revealed the outer membrane lipid phthiocerol dimycocerosate (PDIM) as a key determinant of *M. tuberculosis* antibiotic tolerance triggered by nutrient starvation. Our study implicates PDIM synthesis as a potential target for development of new TB drugs that would sensitize *M. tuberculosis* to existing antibiotics to shorten TB treatment.

## INTRODUCTION

Tuberculosis (TB) infections caused by *Mycobacterium tuberculosis* are notoriously difficult to treat, requiring a lengthy 6-9 month course of combination antibiotic therapy to achieve cure (1). This is due, in part, to *M. tuberculosis* phenotypic antibiotic tolerance that prolongs survival of exposure to drugs. Phenotypic tolerance contributes to the emergence of drug resistance during treatment of other bacterial infections (2, 3) and increases the frequency of drug-resistant *M. tuberculosis* mutants *in vitro* (4). Multidrug-resistant (MDR) *M. tuberculosis*, which are resistant to the two most effective first-line agents rifampicin (RIF) and isoniazid (INH), account for 3-4% of the 10 million newly diagnosed active TB cases annually (5). MDR-TB is more challenging to treat, requiring up to two years of therapy with less effective second-line agents (6). Identifying the molecular mechanisms driving *M. tuberculosis* drug tolerance will be critical to shorten TB treatment and limit emergence of new drug-resistant mutant strains.

Drug efficacy can be reduced by heritable drug resistance mutations, which allow bacteria to grow in the presence of the antibiotic, or by non-heritable phenotypic drug tolerance and persistence, which increase the time required for an antibiotic to kill the bacterial population (7). Tolerance is defined as increased recalcitrance of the entire bacterial population to the drug while antibiotic persisters are a subpopulation of bacteria that survive drug exposure and are typically revealed by biphasic killing in time-based antibiotic kill curve assays (7). Persisters can be either spontaneous, arising in a subset of the population via stochastic mechanisms, or triggered by environmental signals, such as nutrient starvation, stress, or even antibiotics themselves that drive bacteria into a slower growing state (7). The distinction between tolerance and persistence is blurry, but triggers that affect the entire bacterial population (*e.g*. nutrient restriction) were proposed to increase tolerance (8). In *M. tuberculosis*, drug tolerance can arise through any mechanism that decreases target vulnerability or antibiotic interaction with its target (9), including slow growth (10, 11), reduced metabolism (12), decreased intracellular drug concentration (13) or slow pro-drug activation (14).

*M. tuberculosis* persisters exist as a small proportion of exponentially growing cultures *in vitro* (15), but drug tolerance can be triggered by stress. These include stressors encountered during infection such as nutrient starvation, iron limitation, acidic pH, and hypoxia or low oxygen saturation, all of which slow *M. tuberculosis* growth and trigger drug tolerance *in vitro* (15–19). Drug tolerant *M. tuberculosis* have been observed within infected mouse lungs (20), in interferon-gamma (IFN-γ) activated macrophages (21), in the hypoxic caseum of necrotic granulomas (22) and in human sputum (23, 24).

Genetic approaches including identification of genes expressed in the drug-tolerant population (25, 26) and screens for strains exhibiting altered drug tolerance *in vitro* (27, 28) or during infection of macrophages or mice (29–32) have revealed some molecular mechanisms underlying *M. tuberculosis* drug tolerance. Specific metabolic pathways and toxins of toxin-antitoxin (TA) systems induce *M. tuberculosis* drug tolerance by slowing growth. For example, loss of function mutations in the glycerol kinase gene *glpK*, which is required for growth on glycerol that is a primary carbon source in the standard Middlebrook culture medium, increase drug tolerance *in vitro* and during infection (27, 30, 33). Many TA systems are up-regulated in the drug tolerant subpopulation (25, 26) and the toxins generally act to inhibit bacterial growth (34). The VapC12 toxin, a RNase that targets *proT* tRNA, was specifically implicated in drug tolerance of *M. tuberculosis* grown on cholesterol by slowing bacterial replication (35).

Nutrient starvation is one signal that triggers *M. tuberculosis* drug tolerance (16), but the molecular mechanisms underlying this process remain poorly characterized. *M. tuberculosis* growth arrest and tolerance to INH during nutrient starvation requires production of the stringent response alarmone (p)ppGpp by Rel (12), but whether Rel activity is required for tolerance to other antibiotics has not been explored. Nutrient-starved *M. tuberculosis* exhibit lower intracellular concentrations of RIF and fluoroquinolone antibiotics, which may be due to reduced drug uptake as drug efflux pump inhibitors did not reverse the drug tolerance induced by nutrient restriction (36). However, the genetic determinants that promote *M. tuberculosis* drug tolerance triggered by nutrient limitation have not been comprehensively identified.

Here we describe a genetic screen using transposon (Tn) sequencing (Tn-seq) to identify and characterize *M. tuberculosis* factors that influence antibiotic tolerance triggered by nutrient limitation. We identify over 100 *M. tuberculosis* Tn mutants with altered drug tolerance in either phosphate-starved or stationary phase conditions. We confirm decreased antibiotic tolerance phenotypes of individual Tn mutants, but show that for two of these mutants the phenotypes are unlinked to the Tn-disrupted gene. Instead, we find that secondary mutations preventing production of the outer membrane lipid phthiocerol dimycocerosate (PDIM) cause increased susceptibility to antibiotics. As PDIM is also a critical virulence determinant (37), our findings suggest that PDIM synthesis is an attractive target for development of new drugs that would both decrease virulence and sensitize *M. tuberculosis* to existing antibiotics.

## RESULTS

### Construction of an arrayed and sequence-mapped *M. tuberculosis* Erdman transposon mutant library

To identify *M. tuberculosis* determinants of drug tolerance in nutrient starvation, we planned to screen transposon (Tn) mutants for those with defects surviving drug exposure. Since a large percentage of the population is killed by antibiotic treatment, we expected our screen to have an inherent bottleneck that would cause stochastic loss of individual Tn mutant strains. To overcome this bottleneck, we screened defined pools of Tn mutants, which we created as an arrayed library. To enable recovery of auxotrophs, Tn mutants were selected on a nutrient-rich medium, MtbYM, that contains additional carbon and nitrogen sources, vitamins and co-factors compared to the standard Middlebrook 7H9 medium (38). Approximately 8000 *M. tuberculosis* Erdman Tn mutants were arrayed in 80 racks, each with 96 barcoded tubes. Tn mutant pools for experiments were created by combining all ~96 Tn mutants in a rack.

To facilitate recovery of Tn mutants of interest, we used orthogonal pooling and Tn-seq to map the location of Tn mutants in the library. The library was divided into two sets of 40 racks. For each set of 40 racks, pools were created of all mutants in each row (rows A-H, 8 pools, each with 12 x 40 = 480 mutants), all mutants in each column (columns 1-12, 12 pools, each with 8 x 40 = 320 mutants) and all mutants in each rack (racks 1-40 or 41-80, 40 pools, each with 96 mutants) to generate 60 pooled samples per set of racks for Tn-seq. For Tn mutants with no sibling clones in the set of racks, sequence reads corresponding to the Tn insertion site appear in equal abundance in one rack pool, one column pool and one row pool. For racks 1-40, we used two mapping methods: the heuristic Straight Three strategy (39) and the probabilistic Knockout Sudoku algorithm (40). We found good agreement between these mapping methods, with a larger percentage of Tn mutants mapped by Knockout Sudoku (Table S1). Mutants in racks 41-80 were mapped only by Knockout Sudoku (Table S1).

Our arrayed library contains 11,182 total Tn insertions at 6,842 unique locations in the *M. tuberculosis* Erdman genome. These include 1,323 unique insertions in intergenic regions and 5,519 unique insertions within annotated open reading frames (ORFs). We identified Tn insertions in 2,328 of the 3,102 genes previously described as being non-essential for growth of *M. tuberculosis* H37Rv in MtbYM rich medium (~75% coverage) (38). Tn insertions were distributed evenly throughout the *M. tuberculosis* Erdman genome (Fig S1A). We confidently mapped the location of 6,262 Tn mutants (55.97%) by Knockout Sudoku, 3,840 of which were mapped to tubes that contained a single Tn insertion (Table S1; Fig S1B).

### Tn-seq identifies starvation-induced drug tolerance determinants

To identify nutrient-limited conditions that reproducibly increase *M. tuberculosis* antibiotic tolerance, we either grew cultures to stationary phase (a general nutrient limitation) or starved the cultures of inorganic phosphate (P_i_, a defined nutrient limitation). We used combinations of two drugs, each with different modes of action, to prevent emergence of drug-resistant mutants: ciprofloxacin and isoniazid (CIP+INH) or rifampicin and isoniazid (RIF+INH). Each combination included a bactericidal drug (RIF or CIP) and INH at a bacteriostatic low dose to promote isolation of persister variants (15). We compared WT *M. tuberculosis* Erdman drug tolerance in these conditions between Middlebrook 7H9 and MtbYM rich media. The rate at which *M. tuberculosis* was killed by antibiotics was decreased in MtbYM compared to 7H9 medium in both stationary phase and P_i_-starved conditions and with both the CIP+INH and RIF+INH drug combinations (Fig 1A and 1B). These data suggest that the additional nutrient sources in MtbYM promote survival of drug-treated *M. tuberculosis* and that the conditions selected for our screen induce high drug tolerance to optimize recovery of Tn mutants with increased susceptibility to antibiotics.

**Fig 1.**
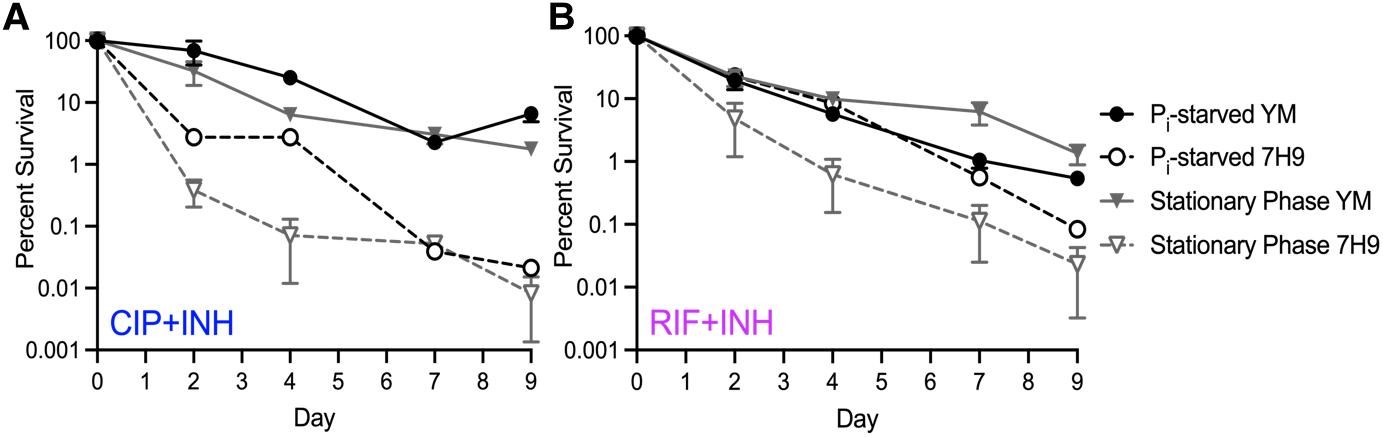
Growth of *M. tuberculosis* in MtbYM rich medium increases starvation-induced antibiotic tolerance. Wild-type *M. tuberculosis* Erdman was grown in 7H9 or MtbYM media to stationary phase or starved of inorganic phosphate (P_i_) for 72 hours before adding the antibiotics ciprofloxacin (CIP, 8 μg/ml) plus isoniazid (INH, 0.2 μg/ml) (**A**) or rifampicin (RIF, 0.1 μg/ml) plus INH (0.2 μg/ml) (**B**). Cultures were incubated at 37°C with aeration for nine days. Drug-tolerant bacteria were enumerated by serially diluting and plating on 7H10 agar at the indicated time points. The mean ± SEM of at least two independent experiments are shown.

To identify *Mtb* determinants of starvation-induced drug tolerance, we screened low-complexity pools of Tn mutants in the starvation and drug treatment conditions we developed and identified mutants with altered fitness by Tn-seq. We note that our screen cannot distinguish between mutants that alter intrinsic drug resistance, drug tolerance, or formation of persister variants, any of which would alter Tn mutant fitness upon drug exposure. Briefly, Tn mutant pools were grown to stationary phase (7 days in MtbYM) or P_i_-starved (72 hours in P_i_-free MtbYM). Each starved culture was plated on MtbYM agar prior to drug exposure as an input control and then split into triplicate no drug control, CIP+INH treated, or RIF+INH treated experimental groups. Cultures were incubated for 9 days before plating on MtbYM agar to recover surviving Tn mutants. Tn mutant abundance in each experimental condition (input, no drug output, CIP+INH output, RIF+INH output) was determined by Tn-seq. In preliminary experiments, we determined that we could screen ~500 Tn mutants (pools of 5 racks) simultaneously without stochastic loss of mutants in individual biological replicates.

We screened two pools, each with ~500 Tn mutants (racks 6-10 and racks 16-20) and obtained similar numbers of Tn-seq reads mapped to the *M. tuberculosis* genome across all experimental conditions (Table S2). To determine the fitness of Tn mutants, we compared the normalized frequency of sequence reads at each Tn insertion site between experimental and either input or no drug control conditions using TnseqDiff, which is compatible with low-density Tn libraries (41). Complete TnseqDiff analyses are available in Table S3. Using ≥ ±2 Log_2_ fold change and adjusted *P*-value < 0.025 statistical significance cut-offs, we identified 122 Tn insertions that exhibited differential fitness in one or more comparisons corresponding to 17 intergenic insertions and 92 unique ORFs disrupted (Table S4).

In the P_i_ starvation condition, we identified 102 Tn mutants with significantly altered fitness (Fig 2, Table S4). Of the 86 Tn mutants with reduced fitness (negative fold change), 11 showed phenotypes in the no drug control/input comparison (Fig 2C), suggesting that these gene products are required for survival of P_i_ starvation. These included Tn insertions in genes putatively involved in nucleotide metabolism or transport (*purN*, *pyrR*, *mkl*), central metabolism (*pckA*), cell division (*ftsX*) and stress responses (*htpX*, *uvrB*) (Fig 2C, Table S4). We identified 76 mutants that showed reduced fitness in the CIP+INH/input comparison (Fig 2A) and 11 mutants with reduced fitness in the RIF+INH/input comparison (Fig 2B). Of these, seven mutants exhibited reduced fitness in both drug treatment conditions (Fig 2D, Table 1). Similar TnseqDiff analyses were done comparing relative Tn mutant abundance in each drug treatment condition to the no drug control. Only four Tn mutants were identified that met our statistical significance criteria in these comparisons (Table S4; Fig S2).

**Fig 2.**
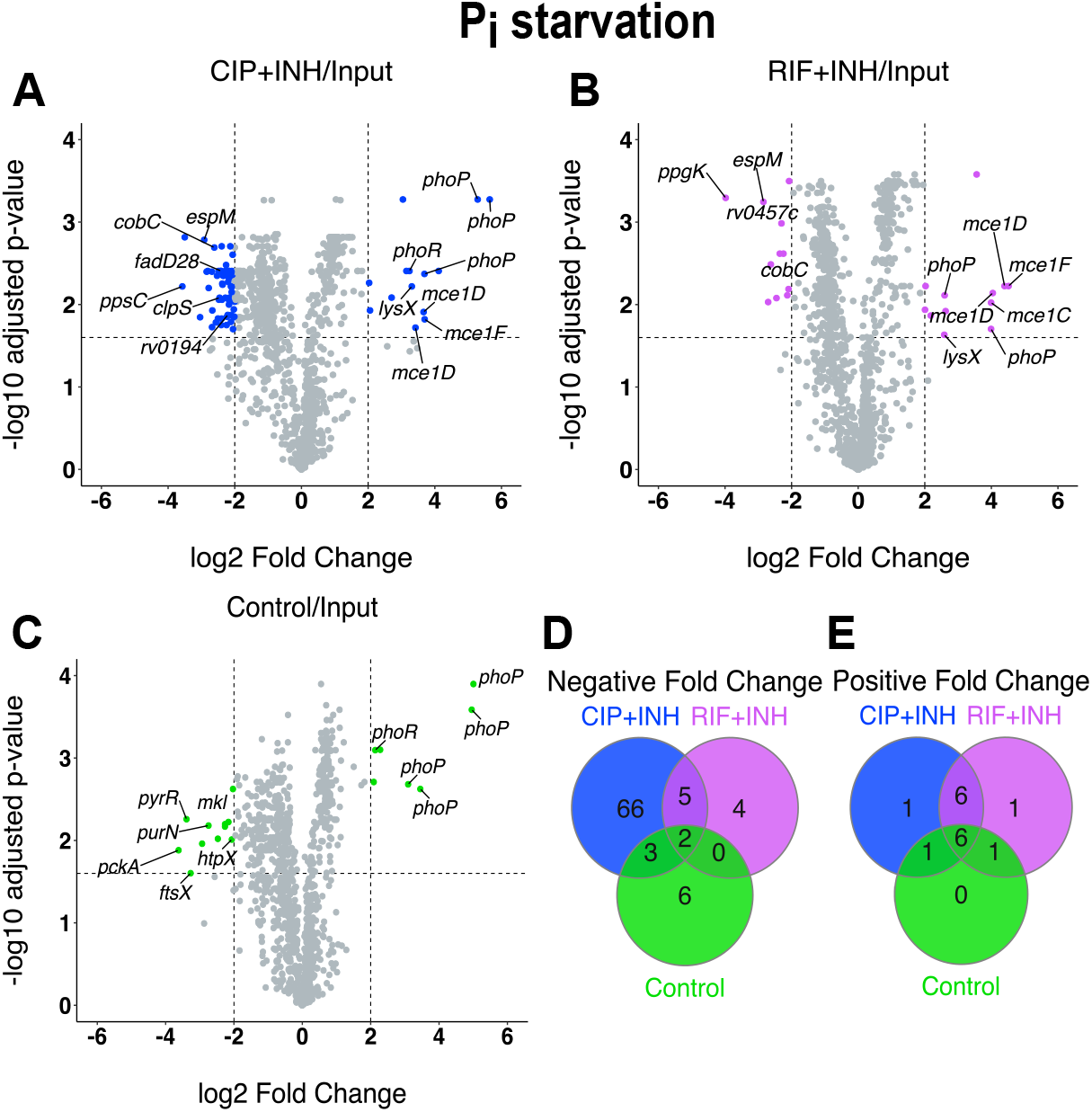
Mutants with altered fitness upon drug treatment in P_i_-limited MtbYM medium. (**A-C**) Volcano plots of TnseqDiff statistical analysis of Tn-seq data for P_i_ starved Tn mutant pools treated with CIP+INH (**A**), RIF+INH (**B**), or no drug control (**C**) compared to input. Dashed lines indicate ± 2 Log_2_ fold change and adjusted *p*-value < 0.025 statistical significance cutoffs. Tn mutants meeting significance are colored. (**D-E**) Venn diagrams displaying the number of Tn mutants with significant negative (**D**) or positive (**E**) fold changes in relative fitness.

**Table 1.** Subset of Tn mutants identified by TnseqDiff as having significantly altered fitness in antibiotic-treated P_i_ starved cultures.^a^

| Position<sup>c</sup> | Locus tag | Gene | Gene product function | Log<sub>2</sub> fold change (adjusted *P* value)<sup>b</sup> |  |  |
| --- | --- | --- | --- | --- | --- | --- |
|  |  |  |  | Control/Input | CIP+INH/Input | RIF+INH/Input |
| 226628 | Erdman_0220 | *rv0194* | Drug transporter | -1.33 (0.013) | **-2.21 (0.013)** | -1.04 (0.006) |
| 229774 | Erdman_0220 | *rv0194* | Drug transporter | -1.15 (0.06) | **-2.25 (0.018)** | -0.94 (0.046) |
| 2496685 | Erdman_2452 | *cobC* | Aminotransferase, cobalamin synth | -1.43 (0.011) | **-2.61 (0.002)** | **-2.25 (0.002)** |
| 3247032 | Erdman_3215 | *ppsC* | PDIM synthesis | -1.78 (0.008) | **-3.57 (0.006)** | -0.628 (0.12) |
| 3273492 | Erdman_3223 | *fadD28* | PDIM synthesis | -1.29 (0.009) | **-2.41 (0.004)** | -0.81 (0.022) |
| 3395716 | Erdman_3338 | *rv3049c* | Mono-oxygenase | -1.27 (0.009) | **-2.01 (0.005)** | -0.76 (0.03) |
| 3397057 | Erdman_3338 | *rv3049c* | Mono-oxygenase | -1.57 (0.008) | **-2.17 (0.005)** | -0.96 (0.006) |
| 4320034 | Erdman_4236 | *espM* | Transcriptional regulator of ESX-1 | -1.81 (0.009) | **-2.92 (0.002)** | **-2.85 (0.0006)** |
| 201398 | Erdman_0196 | *mce1C* | Fatty acid transport | -0.25 (0.362) | **3.37 (0.028)** | **3.99 (0.009)** |
| 203040 | Erdman_0197 | *mce1D* | Fatty acid transport | -0.28 (0.019) | **3.66 (0.012)** | **4.40 (0.006)** |
| 203224 | Erdman_0197 | *mce1D* | Fatty acid transport | -0.13 (0.409) | **3.42 (0.019)** | **4.04 (0.007)** |
| 205440 | Erdman_0199 | *mce1F* | Fatty acid transport | -0.10 (0.389) | **3.69 (0.015)** | **4.51 (0.006)** |
| 1833015 | Erdman_1805 | *lysX* | Lysinylation of peptidoglycan | 1.82 (0.002) | **3.31 (0.006)** | **2.59 (0.023)** |
| 3831881 | Erdman_3755 | *rv3430c* | Transposase | 0.33 (0.142) | **2.06 (0.012)** | **2.63 (0.012)** |

- This table includes only Tn insertions within open reading frames for which either multiple independent Tn insertions with significant phenotypes were observed in the same gene or pathway or for which significant phenotypes were observed in both drug treatment conditions, but not the no drug control. - Log<sub>2</sub> fold change and adjusted *P* values were determined using TnseqDiff relative to the input control. Bolded text indicates comparisons for which the Tn mutant met the > ± 2 Log<sub>2</sub> fold change and adjusted *P* value < 0.025 statistical significance cutoffs. - First nucleotide position of the Tn insertion site in the *M. tuberculosis* Erdman ATCC 35801 genome AP012340.1.

Mutants significantly impaired for survival of CIP+INH treatment during P_i_ starvation included two with Tn insertions in genes required for production of the outer membrane lipid phthiocerol dimycocerosate (PDIM; *ppsC*, *fadD28*; Table 1). PDIM limits permeability of the *M. tuberculosis* outer membrane to small hydrophilic nutrients including glucose and glycerol (42, 43) and may also restrict diffusion of certain antibiotics, such as the glycopeptide vancomycin (44). We also identified two independent Tn insertions each in Erdman_0220 (*rv0194*), which encodes a ATP-binding cassette (ABC) type efflux pump previously implicated in intrinsic resistance of *Mycobacterium bovis* BCG to multiple drugs, including ampicillin (45), and Erdman_3338 (*rv3049*), which encodes a putative mono-oxygenase (Fig 2A, Table 1).

In P_i_ starvation, we also identified 16 mutants with significantly increased fitness (positive fold change). These included five independent Tn insertions in *phoPR*, which encodes a two-component system that responds to acid stress (46–48). Although the *phoPR* mutants exhibited increased fitness in both drug combinations (Fig 2A and 2B), they were also more abundant in the no drug control/input comparison (Fig 2C), suggesting that PhoPR normally impairs survival of P_i_ limitation, rather than specifically altering antibiotic susceptibility. We identified eight additional Tn mutants with increased abundance in both the CIP+INH/input and RIF+INH/input comparisons (Fig 2A, 2B, 2E). These included four independent Tn insertions in the *mce1* locus (Fig 2A and 2B, Table 1), which encodes a fatty acid transporter (49), suggesting Mce1 contributes to drug susceptibility.

In the stationary phase condition, we identified 32 Tn mutants with significantly altered fitness. In comparisons with the input control, three Tn mutants decreased in relative abundance (negative fold change) and 25 Tn mutants increased in relative abundance (positive fold change) (Fig 3, Table S4). Only two Tn mutants (*pe_pgrs31*::Tn and an intergenic insertion 5’ of *rv3796*) exhibited decreased fitness in drug-treated stationary phase cultures (Fig 3B). In comparisons of drug-treated cultures to the no drug control, Tn mutants in *phoP* and Erdman_3938 (*rv3553*) showed significantly decreased abundance (Fig S3, Table S4). However, these mutants also increased in relative abundance in the no drug control/input comparison (Fig 3C), suggesting that these mutants have a survival advantage in stationary phase.

**Fig 3.**
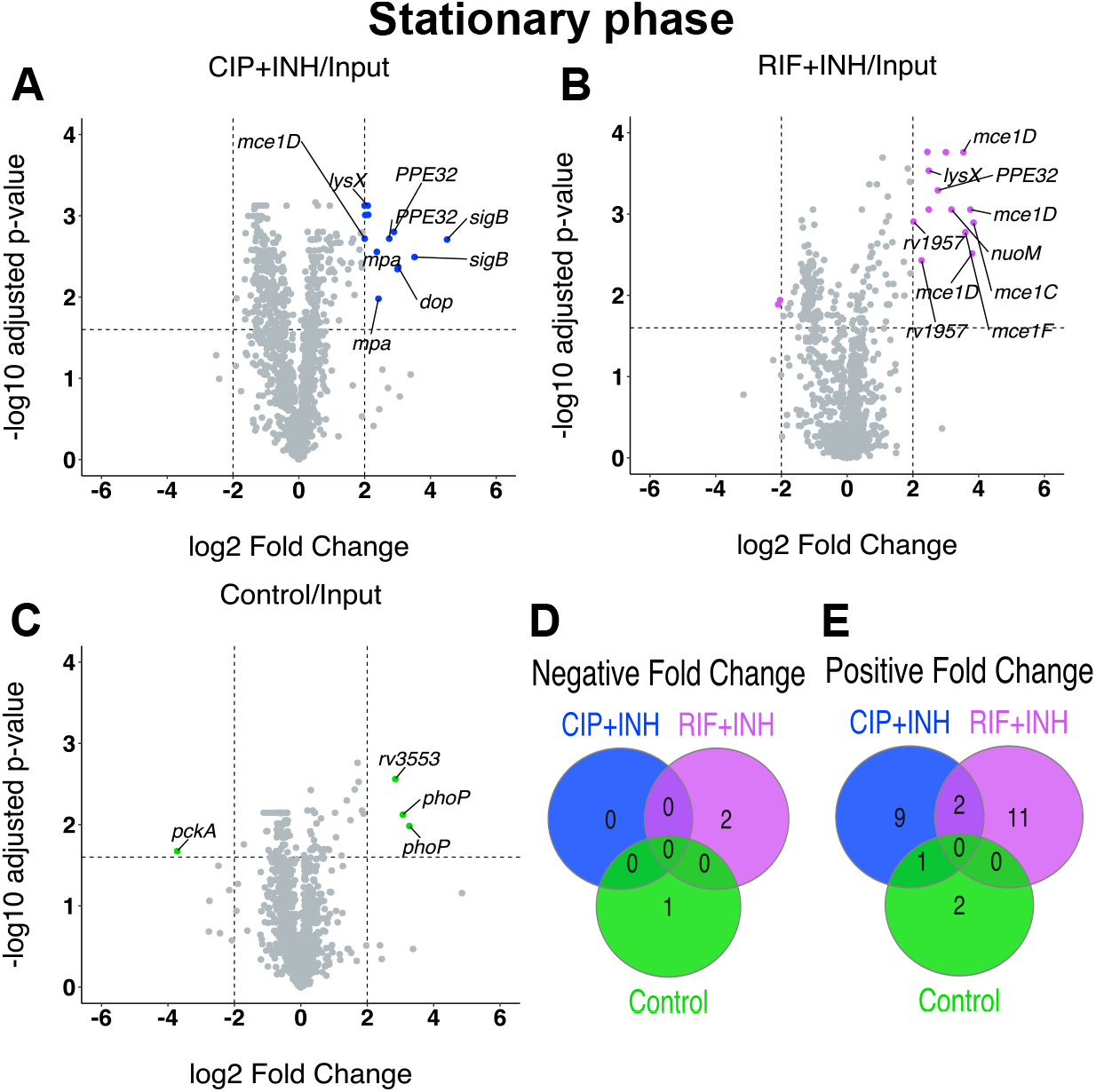
Mutants with altered fitness upon drug treatment during stationary phase in MtbYM medium. (**A-C**) Volcano plots of TnseqDiff statistical analysis of Tn-seq data for Tn mutant pools grown to stationary phase in MtbYM and treated with CIP+INH (**A**), RIF+INH (**B**), or no drug control (**C**) compared to input. Dashed lines indicate ± 2 Log_2_ fold change and adjusted *p*-value < 0.025 statistical significance cutoffs. Tn mutants meeting significance are colored. (**D-E**) Venn diagrams displaying the number of Tn mutants with significant negative (**D**) or positive (**E**) fold changes in relative fitness.

Among the Tn insertions with increased fitness in drug-treated stationary phase cultures, we identified five independent Tn insertions the *mce1* locus (Fig 3A and 3B, Table 2). The *mce1* mutants were all over-represented in RIF+INH-treated treated cultures compared to either the input or no drug controls (Fig 3A and 3B, Fig S3). We also identified multiple independent Tn insertions in *mpa*, *sigB*, the *nuo* operon, *ppe32*, and Erdman_2155 (*rv1957*) that caused increased fitness in drug-treated stationary phase cultures (Fig 3A and 3B, Table 2). Rv1957 is a SecB-like chaperone of the antitoxin HigA1 (50). In the absence of Rv1957, HigA1 is degraded by the ClpXP protease (51), freeing the HigB1 toxin to degrade mRNA and tmRNA (52). Mutation of *rv1957* is expected to limit bacterial replication via HigB1 toxin activation, thereby enhancing drug tolerance. Mpa is the ATPase of the mycobacterial proteasome, which degrades proteins that are post-translationally modified with the prokaryotic ubiquitin-like protein Pup (53). While the proteasome itself has not been directly implicated in mycobacterial drug tolerance, several toxins and antitoxins of TA systems are modified by Pup and may be stabilized in mutants lacking proteasome activity (54, 55).

**Table 2.** Subset of Tn mutants identified by TnseqDiff as having significantly altered fitness in antibiotic-treated stationary phase cultures.^a^

| Position<sup>c</sup> | Locus tag | Gene | Gene product function | Log<sub>2</sub> fold change (adjusted *P* value)<sup>b</sup> |  |  |
| --- | --- | --- | --- | --- | --- | --- |
|  |  |  |  | Control/Input | CIP+INH/Input | RIF+INH/Input |
| 201398 | Erdman_0196 | *mce1C* | Fatty acid transport | 0.23 (0.61) | 1.50 (0.005) | **2.84 (0.001)** |
| 201967 | Erdman_0197 | *mce1D* | Fatty acid transport | -0.26 (0.20) | **2.00 (0.002)** | **3.80 (0.003)** |
| 203040 | Erdman_0197 | *mce1D* | Fatty acid transport | -0.08 (0.59) | 1.26 (0.004) | **2.75 (0.009)** |
| 203244 | Erdman_0197 | *mce1D* | Fatty acid transport | -0.10 (0.38) | 1.15 (0.002) | **3.54 (0.0002)** |
| 205440 | Erdman_0199 | *mce1F* | Fatty acid transport | -0.29 (0.22) | 1.20 (0.002) | **3.60 (0.002)** |
| 1833015 | Erdman_1805 | *lysX* | Lysinylation of peptidoglycan | 1.03 (0.007) | **2.09 (0.0008)** | **2.48 (0.0003)** |
| 2041171 | Erdman_1997 | PPE32 | Unknown PPE family protein | 0.23 (0.03) | 1.13 (0.002) | **2.76 (0.0005)** |
| 2041579 | Erdman_1997 | PPE32 | Unknown PPE family protein | 1.62 (0.004) | **2.75 (0.002)** | 0.68 (0.20) |
| 2041961 | Erdman_1997 | PPE32 | Unknown PPE family protein | 1.74 (0.003) | **2.89 (0.002)** | 0.63 (0.51) |
| 2194032 | Erdman_2155 | *rv1957* | TA chaperone | 0.27 (0.23) | 1.51 (0.007) | **2.01 (0.001)** |
| 2194075 | Erdman_2155 | *rv1957* | TA chaperone | 0.51 (0.10) | 1.65 (0.002) | **2.27 (0.004)** |
| 2364483 | Erdman_2328 | *dop* | Deamidase of pup | 1.38 (0.005) | **3.02 (0.004)** | 0.73 (0.49) |
| 2367722 | Erdman_2331 | *mpa* | Proteasome ATPase | 0.56 (0.35) | **2.37 (0.003)** | 0.87 (0.40) |
| 2369014 | Erdman_2331 | *mpa* | Proteasome ATPase | 0.48 (0.39) | **2.42 (0.01)** | 0.84 (0.40) |
| 3009590 | Erdman_2972 | *sigB* | Sigma factor SigB | 0.87 (0.13) | **3.52 (0.003)** | -0.12 (0.87) |
| 3009923 | Erdman_2972 | *sigB* | Sigma factor SigB | 0.39 (0.59) | **4.50 (0.002)** | 0.08 (0.96) |
| 3511066 | Erdman_3458 | *nuoM* | NADH dehydrogenase I | 0.23 (0.42) | 1.60 (0.004) | **3.17 (0.0009)** |
| 3513262 | Erdman_3459 | *nuoN* | NADH dehydrogenase I | 0.85 (0.02) | **2.11 (0.001)** | 0.94 (0.30) |

- This table includes only Tn insertions within open reading frames for which either multiple independent Tn insertions with significant phenotypes were observed in the same gene or pathway or for which significant phenotypes were observed in both drug treatment conditions, but not the no drug control. - Log<sub>2</sub> fold change and adjusted *P* values were determined using TnseqDiff relative to the input control. Bolded text indicates comparisons for which the Tn mutant met the $> \pm 2$ Log<sub>2</sub> fold change and adjusted *P* value $< 0.025$ statistical significance cutoffs. - First nucleotide position of the Tn insertion site in the *M. tuberculosis* Erdman ATCC 35801 genome AP012340.1.

### Retesting confirms increased drug susceptibility of *ppgK*::Tn and *clpS*::Tn mutants

To validate the phenotypes observed in our screen, we determined the sensitivity of individual Tn mutants to antibiotics in monoculture. We selected only Tn mutants with negative fold changes in the TnseqDiff analysis to characterize pathways that, when inhibited, would sensitize *M. tuberculosis* to existing antibiotics. We focused on genes that were not previously implicated in mycobacterial drug tolerance. The three Tn mutants we selected had relatively severe phenotypes based on the TnseqDiff fold change, had the Tn insertion in the middle of the ORF, and were identified only in the P_i_ starvation screen (Table 3). *rv0457c*::Tn and *ppgK*::Tn were also the only two mutants that exhibited significantly reduced fitness in the RIF+INH-treated vs no drug control comparison (Fig S2, Table S4). Each mutant was recovered from the arrayed library and the Tn insertion site was confirmed by PCR and sequencing.

**Table 3.** Tn mutants with significant antibiotic tolerance defects in P_i_ starvation selected for individual retesting.

| Position<sup>b</sup> | Locus tag | Gene | Gene product function | Log<sub>2</sub> fold change (adjusted *P* value)<sup>a</sup> |  |  |
| --- | --- | --- | --- | --- | --- | --- |
|  |  |  |  | Control/Input | CIP+INH/Input | RIF+INH/Input |
| 551129 | Erdman_0502 | *rv0457c* | Probable peptidase | 0.30 (0.06) | -0.36 (0.12) | **-2.31 (0.001)** |
| 201967 | Erdman_1491 | *clpS* | ClpCP protease adaptor | -1.89 (0.0017) | **-2.46 (0.0083)** | -1.55 (0.0072) |
| 3004201 | Erdman_2965 | *ppgK* | Polyphosphate glucokinase | -1.84 (0.002) | -1.99 (0.002) | **-3.98 (0.0005)** |

- a. Log<sub>2</sub> fold change and adjusted *P* values were determined using TnseqDiff relative to the input control. Bolded text indicates comparisons for which the Tn mutant met the > ± 2 Log<sub>2</sub> fold change and adjusted *P* value < 0.025 statistical significance cutoffs. - b. First nucleotide position of the Tn insertion site in the *M. tuberculosis* Erdman ATCC 35801 genome AP012340.1.

The *rv0457c*::Tn mutant exhibited specific susceptibility to RIF+INH in the P_i_ starvation screen (Table 3). *rv0457c* encodes a prolyl oligopeptidase (56) and is located immediately 5’ of the *mazE1-mazF1* operon that encodes a TA system. MazF toxins were previously implicated in *M. tuberculosis* drug tolerance (57). The *rv0457c::Tn* mutant displayed a subtle but statistically significant increase in sensitivity to RIF+INH, but not CIP+INH, in P_i_ starvation conditions (Fig 4A and 4B). As the *rv0457c::Tn* mutant did not exhibit strong phenotypes upon retesting, it was not pursued further.

**Fig 4.**
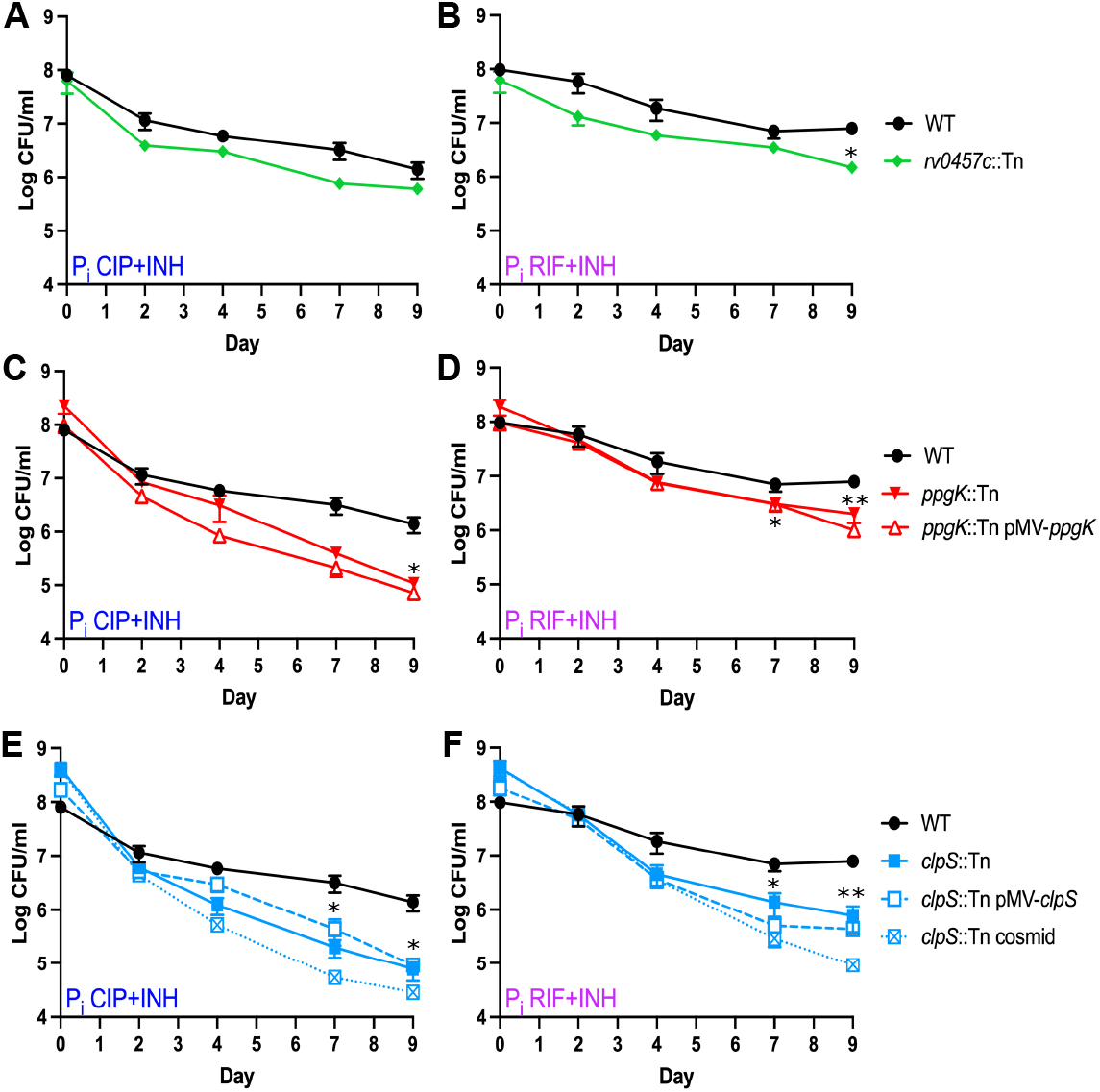
Individual Tn mutants identified in the P_i_ starvation Tn-seq screen exhibit reduced fitness during drug treatment. *M. tuberculosis* strains were P_i_ starved for 72 hr in P_i_-free MtbYM medium before addition the drug combinations ciprofloxacin (CIP, 8 μg/ml) plus isoniazid (INH, 0.2 μg/ml) (**A, C, E**) or rifampicin (RIF, 0.1 μg/ml) plus INH (0.2 μg/ml) (**B, D, F**). Surviving bacteria were enumerated by plating serial dilutions on 7H10 agar. Data represent the average ± SEM of at least two independent experiments. Asterisks indicate statistically significant differences between Tn mutant and WT: * *P* < 0.05, ** *P* <0.005.

The *ppgK*::Tn mutant exhibited the highest sensitivity to RIF+INH in P_i_ starved conditions in our screen (Fig 2B) and was also susceptible to the CIP+INH combination, though it did not reach our statistical significance cutoffs (Table 3). *ppgK* encodes the dominant glucokinase in *M. tuberculosis* (58), catalyzing phosphorylation of glucose with a preference for polyphosphate as the phosphodonor (59). The *ppgK*::Tn mutant was significantly more susceptible to both CIP+INH and RIF+INH in P_i_-free MtbYM medium (Fig 4C and 4D). However, the minimal inhibitory concentration (MIC_90_) for all three drugs was similar between the *ppgK*::Tn and WT strains, suggesting that the *ppgK*::Tn mutant has altered antibiotic tolerance (Table 4). We attempted to complement these phenotypes by providing *ppgK in trans* using a construct similar to that previously reported to complement a Δ*ppgK* mutant (58). Quantitative RT-PCR (qRT-PCR) confirmed *ppgK* transcription from the pMV-*ppgK* vector (data not shown), but complementation did not increase the tolerance of the *ppgK*::Tn mutant to either drug combination (Fig 4C and 4D). These data suggest that the *ppgK*::Tn mutant harbors a secondary mutation, unlinked to the Tn, that causes increased susceptibility to antibiotics.

**Table 4.**
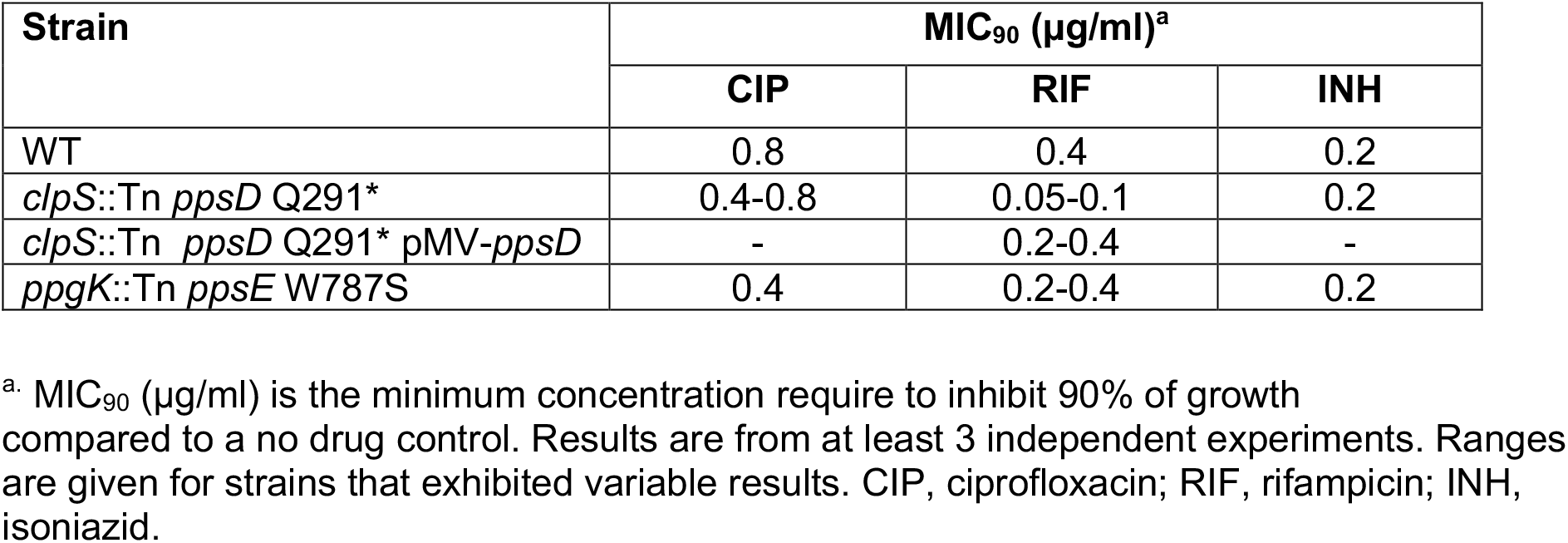
Minimum inhibitory concentrations of antibiotics against *M. tuberculosis* grown in MtbYM rich medium.

The *clpS*::Tn (*rv1331*::Tn) mutant exhibited high susceptibility to CIP+INH, but not RIF+INH, in P_i_-starved conditions in our screen (Table 3). *clpS* encodes an adaptor for the *M. tuberculosis* ClpC1P1P2 (ClpCP) protease. ClpS promotes ClpCP degradation of proteins with destabilizing N-terminal residues (the N-end rule) and inhibits degradation of SsrA-tagged proteins derived from translationally stalled ribosomes (60, 61). *M. tuberculosis* ClpCP was implicated in drug tolerance because it degrades antitoxins from several classes of TA systems (61). Loss of the ClpS adaptor may therefore alter the stability of certain ClpCP protease substrates that influence drug tolerance. When tested in monoculture, the *clpS*::Tn mutant was highly susceptible to both CIP+INH and RIF+INH in P_i_-free MtbYM medium (Fig 4E and 4F). While the *clpS*::Tn mutant had a similar MIC_90_ for both CIP and INH relative to the WT control, the MIC_90_ for RIF was reduced 4-to 8-fold, suggesting that the *clpS*::Tn mutant has reduced intrinsic resistance to RIF (Table 4). We attempted to complement the *clpS*::Tn mutant by providing *clpS in trans*. We observed *clpS* transcript from pMV-*clpS* by qRT-PCR (data not shown), but the complemented strain remained susceptible to both drug combinations (Fig 4E and 4F). As *clpS* is encoded at the 5’ end of a putative operon, we considered the possibility that the Tn insertion was polar on expression of downstream genes. A cosmid covering the complete *clpS* region also failed to complement the *clpS*::Tn mutant phenotype (Fig 4E and 4F). These data suggest that the *clpS*::Tn mutant also has a secondary mutation that increases susceptibility to antibiotics.

### Tn mutants with defects in drug tolerance harbor secondary mutations that disrupt production of phthiocerol dimycocerosate (PDIM) and cause decreased drug tolerance

Since neither the *ppgK*::Tn nor *clpS*::Tn mutant phenotypes could be complemented, we sought to identify secondary mutations responsible for their drug susceptibility phenotypes. We conducted whole genome resequencing on *rv0457c*::Tn, *ppgK*::Tn, *clpS*::Tn and our WT *M. tuberculosis* Erdman strain. This sequencing confirmed the predicted Tn insertion sites in each strain, and demonstrated that each strain harbored a single Tn, ruling out the possibility that a secondary Tn insertion was responsible for their phenotypes (Fig S4A-C). In both the *ppgK*::Tn and *clpS*::Tn mutants, we identified non-synonymous single nucleotide polymorphisms (SNPs) in genes required for production of the lipid phthiocerol dimycocerosate (PDIM). No mutations in genes required for PDIM biosynthesis were identified in the WT Erdman or *rv0457c*::Tn strains. The *ppgK*::Tn strain had a G to C mutation at position 2360 in *ppsE* (Fig S4D), which encodes a polyketide synthase required for production of the phthiocerol chain of PDIM (62). This SNP is predicted to cause a W787S amino acid substitution in PpsE that may alter PpsE activity and PDIM production. The *clpS*::Tn strain had a C to T mutation at position 655 in *ppsD* (Fig S4E), which is predicted to introduce a premature amber stop codon at position 219 in PpsD. As *ppsD* also encodes a polyketide synthase required for production the phthiocerol component of PDIM (62), the *ppsD* Q219* mutation is predicted to completely block PDIM production. Excluding highly repetitive sequences, such as the PE and PPE genes, which are difficult to resolve by short-read sequencing, these were the only non-synonymous SNPs identified in the *ppgK*::Tn and *clpS*::Tn strains.

To directly test whether the *ppsD* Q219* or *ppsE* W787S mutations block PDIM production by the *clpS*::Tn or *ppgK*::Tn strains, respectively, we analyzed PDIM production by an established radiolabeling method. Bacteria were labeled with ^14^C propionate, which is selectively incorporated into PDIM, and the PDIM (DIM A) and phthiodiolone dimycocerosate precursor (DIM B) were detected in apolar lipid extracts by thin layer chromatography (37, 63). As expected, the *clpS*::Tn *ppsD* Q219* mutant did not produce any detectable PDIM (Fig 5A, lane 2). The *ppgK*::Tn *ppsE* W787S mutant exhibited an intermediate PDIM production phenotype, with a 2.3-fold reduction in both DIM A and DIM B compared to the WT control (Fig 5A, lane 4). These results suggest that the antibiotic susceptibility phenotypes of both mutants could be due to reduced PDIM production rather than the Tn insertion. These data also suggest that the intermediate drug susceptibility phenotypes of the *ppgK*::Tn *ppsE* W787S mutant could be caused by its intermediate level of PDIM production.

**Fig 5.**
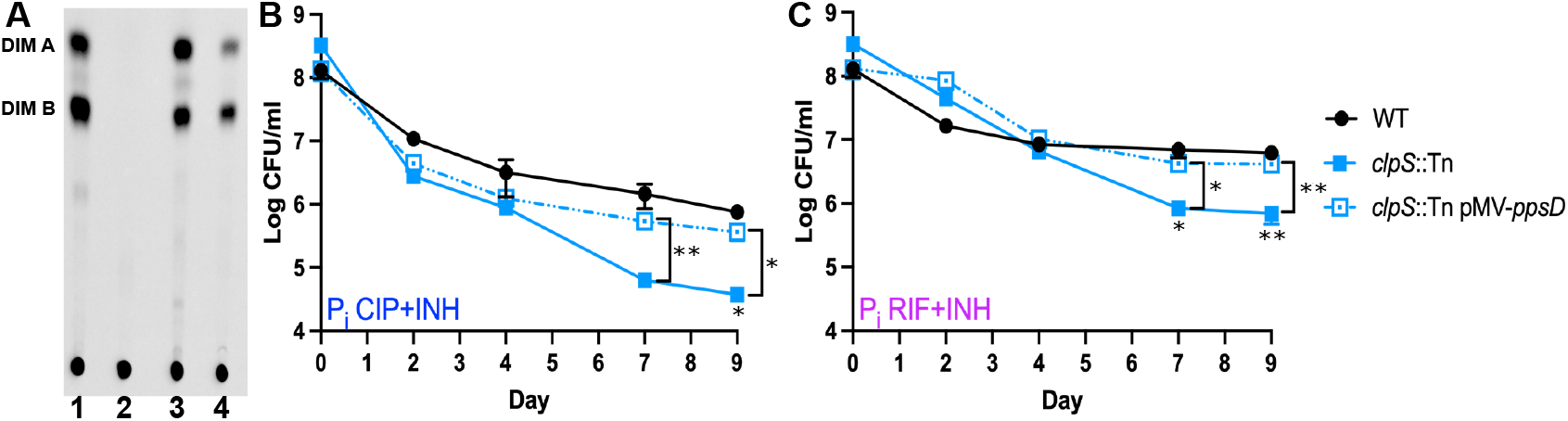
Loss of PDIM causes increased drug susceptibility of the *clpS*::Tn mutant. **A**) Thin-layer chromatographic analysis of PDIM lipids extracted from [^14^C] propionate-labeled WT Erdman (lane 1), *clpS*::Tn *ppsD* Q291* (lane 2), *clpS*::Tn pMV-*ppsD* (lane 3), and *ppgK*::Tn *ppsE* W787S (lane 4). (**B-C**) *M. tuberculosis* strains were P_i_ starved for 72 hr in P_i_-free MtbYM medium before adding ciprofloxacin (CIP, 8 μg/ml) plus isoniazid (INH, 0.2 μg/ml) (**B**) or rifampicin (RIF, 0.1 μg/ml) plus INH (0.2 μg/ml) (**C**). Surviving bacteria were enumerated by plating serial dilutions on 7H10 agar. Data represent the average ± SEM of three biological replicates. Asterisks indicate statistically significant differences between WT and *clpS*::Tn (below points) or between *clpS*::Tn and *clpS*::Tn pMV-*ppsD* (brackets): * *P* < 0.05, ** *P* <0.005.

To determine if PDIM deficiency caused increased susceptibility of the *clpS*::Tn *ppsD* Q219* mutant to antibiotics, we complemented the *ppsD* Q219* mutation with *ppsD* on a plasmid. Complementation with *ppsD* fully restored PDIM production (Fig 5A, lane 3). We tested the sensitivity of the *ppsD* complemented strain to both CIP+INH and RIF+INH in P_i_ starvation conditions and observed similar resistance to both drug combinations as the WT control (Fig 5B and 5C). These data demonstrate that loss of PDIM production, rather than loss of ClpS function, causes increased drug susceptibility of the *clpS*::Tn *ppsD* Q219* mutant.

### PDIM-deficient mutants are hypersusceptible to antibiotics in stationary phase and exponential phase growth conditions

Although the *clpS*::Tn *ppsD* Q219* and *ppgK*::Tn *ppsE* W787S mutants were initially identified only in the P_i_-starved drug screen, we sought to determine if hypersusceptibility to antibiotics was specific to this growth condition. We therefore tested susceptibility of both mutants to the CIP+INH and RIF+INH drug combinations in stationary phase and exponential phase cultures grown in MtbYM rich medium. In stationary phase, both the *clpS*::Tn *ppsD* Q219* and *ppgK*::Tn *ppsE* W787S mutants showed a significant decrease in tolerance to CIP+INH (Fig 6A). Both mutants also exhibited decreased tolerance to RIF+INH in stationary phase, though this did not achieve statistical significance (Fig. 6B). These data suggest that some Tn mutants with antibiotic tolerance phenotypes were not uncovered by our stationary phase Tn-seq screen, perhaps due to the stringent statistical significance cutoffs we used. The *clpS*::Tn *ppsD* Q219* mutant also exhibited a modest, but statistically significant, decrease in antibiotic tolerance in exponential phase for both CIP+INH and RIF+INH (Fig 6C and 6D). These data suggest that in addition to a role in drug tolerance triggered by starvation, the *clpS*::Tn *ppsD* Q219* mutant also produces fewer stochastic persister variants.

**Fig 6.**
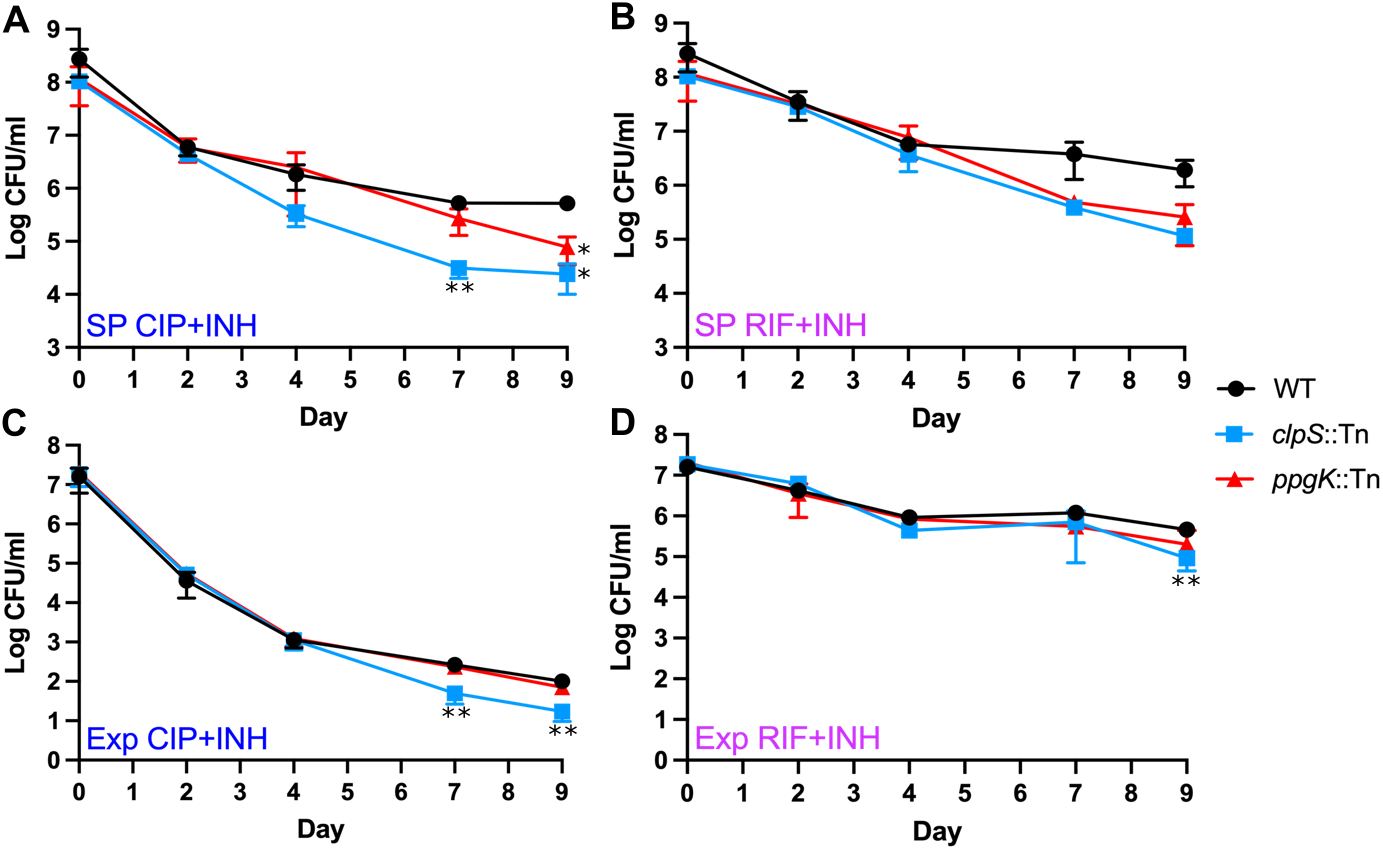
PDIM-deficient mutants are hypersusceptible to antibiotics in stationary and exponential phase MtbYM cultures. *M. tuberculosis* strains were grown to early stationary phase (**A-B**) or exponential phase (**C-D**) in MtbYM medium before adding ciprofloxacin (CIP, 8 μg/ml) plus isoniazid (INH, 0.2 μg/ml) (**A, C**) or rifampicin (RIF, 0.1 μg/ml) plus INH (0.2 μg/ml) (**B, D**). Data represent the average ± SEM of at least two independent experiments. Asterisks indicate statistically significant differences between Tn mutant and WT: * *P* < 0.05, ** *P* <0.005.

## DISCUSSION

Molecular mechanisms driving *M. tuberculosis* recalcitrance to antibiotics under nutrient starvation are poorly characterized. Here, using a Tn-seq screen, we identify PDIM production as a critical determinant of *M. tuberculosis* drug tolerance in nutrient-limited conditions. We identified two Tn mutants, *clpS*::Tn and *ppgK*::Tn, that were hypersusceptible to antibiotics. Both mutants harbored secondary mutations, unlinked to the Tn, that disrupted PDIM production. We restored PDIM production to the *clpS*::Tn strain by complementing the *ppsD* Q291* mutation, and showed that this restored normal drug tolerance, directly demonstrating that *M. tuberculosis* requires PDIM to tolerate antibiotic exposure. Loss of PDIM caused a decrease in the MIC_90_ for RIF, demonstrating that PDIM contributes to intrinsic RIF resistance. However, PDIM-deficient strains also exhibited increased susceptibility to the CIP+INH combination despite no change in the MIC_90_ for these drugs, indicating that PDIM also promotes antibiotic tolerance. The *ppgK*::Tn *ppsE* W787S mutant exhibited an intermediate drug tolerance phenotype that was associated with reduced, but not absent, PDIM production, suggesting that even partial inhibition of PDIM synthesis can sensitize *M. tuberculosis* to antibiotics.

There are at least two mechanisms by which PDIM could increase *M. tuberculosis* drug tolerance: decreasing the intracellular concentration of antibiotics by decreasing permeability of the outer membrane or altering the intracellular concentrations of central metabolites by functioning as a metabolic sink for propionate. PDIM decreases permeability of the *M. tuberculosis* outer membrane to small molecules including glucose and glycerol (42, 43) and may also restrict diffusion of some antibiotics. Indeed, PDIM is required for intrinsic resistance to vancomycin, likely by decreasing vancomycin access to its peptidoglycan target (44). PDIM has previously been implicated in drug tolerance in other mycobacteria. PDIM-deficient *Mycobacterium bovis* BCG exhibited increased susceptibility to RIF, with a 4-fold decrease in MIC, but there was no change in susceptibility to INH or CIP (44). PDIM also increases antibiotic tolerance of *Mycobacterium marinum*, which was correlated with reduced outer membrane permeability (64, 65). We observed that PDIM enhances *M. tuberculosis* drug tolerance particularly in nutrient-limited conditions that limit accumulation of RIF and fluoroquinolone antibiotics (36). Since efflux pump inhibitors did not reverse the drug tolerance triggered by nutrient starvation (36), it is tempting to speculate that PDIM decreases drug uptake in nutrient limited conditions by limiting import of antibiotics through the outer membrane.

Alternatively, PDIM could alter drug tolerance by effects on central metabolism. Synthesis of the long chain branched fatty acids in PDIM requires the metabolite methylmalonyl-CoA, which is derived from propionate (62). During infection, *M. tuberculosis* catabolizes fatty acids and cholesterol, which serve as primary carbon sources, to propionate (66, 67). Excess propionate stimulates increased production of PDIM with longer mycocerosic acid chain lengths both *in vitro* and during infection of macrophages or mice (66, 68). PDIM was therefore proposed to act as a sink for propionyl CoA, which can be toxic at high concentrations (66, 69). PDIM-deficient strains may be more susceptible to antibiotics, particularly in growth conditions with fatty acids or cholesterol as carbon sources, due to the combined effects of the antibiotic and accumulation of toxic central metabolites.

Connections between *M. tuberculosis* central metabolism, production of outer membrane lipids and antibiotic tolerance have previously been reported. *M. tuberculosis* isocitrate lyase (ICL) is required for catabolism of both even- and odd-chain fatty acids and for tolerance to several different classes of antibiotics (70). The increased susceptibility of mutants lacking ICL activity was correlated to accumulation of TCA cycle intermediates and to increased endogenous oxidative stress (70). However, ICL is also required for propionate catabolism (69). Accumulation of toxic propionate metabolites could therefore similarly cause hypersusceptiblity of *icl* mutants to antibiotics. Growth in medium with either propionate or cholesterol as a carbon source also increases intrinsic resistance of *M. tuberculosis* to RIF (71). This increased RIF resistance was correlated with increased production and chain length of sulfolipid-1 (SL-1) (71), another branched-chain outer membrane lipid synthesized from propionate (68). PDIM production and chain length also increase with propionate as a carbon source, and PDIM is much more abundant compared to SL-1 (68), suggesting that PDIM could be primarily responsible for the carbon source dependent increase in RIF resistance.

Our results contrast with a previous study, in which selection for *M. tuberculosis* mutants with higher antibiotic persistence revealed multiple strains harboring spontaneous mutations in genes required for PDIM production (27). PDIM-deficient mutants exhibited increased tolerance to multiple classes of antibiotics in exponential phase in the standard Middlebrook 7H9 medium, which contains glucose and glycerol as primary carbon sources (27). We may have observed decreased antibiotic tolerance of PDIM-deficient strains due to our use of the MtbYM rich medium, which contains branched-chain amino acids and pyruvate that are catabolized to propionate and vitamin B12 that activates production of methylmalonyl-CoA that is used for PDIM synthesis (72, 73). It is unclear which *in vitro* growth medium more closely reflects the conditions *M. tuberculosis* experiences in the host or whether loss of PDIM would enhance drug susceptibility during lung infection. This question will be challenging to address because PDIM is also a critical *M. tuberculosis* virulence determinant that is required for resistance to innate immunity (37, 63, 74). We intend to explore the role of PDIM in antibiotic tolerance during lung infection in our future studies.

Our screen identified over 100 unique *M. tuberculosis* Tn insertion mutants with altered drug tolerance phenotypes, including several genes or pathways with multiple independent Tn insertions. Our results point to the importance of regulated protein degradation in *M. tuberculosis* drug tolerance. Loss of Rv1957, a chaperone of the HigA1 antitoxin, or loss of proteasome components (Mpa or Paf) caused increased drug tolerance, possibly due to stabilization of toxins that inhibit bacterial replication. Mutations in genes encoding the Mce1 system, which is required for uptake of fatty acids (49), also increased antibiotic tolerance. Mce1 may also function in uptake of antibiotics, such that loss of Mce1 reduces antibiotic import. Alternatively, loss of Mce1 function may reduce accumulation of fatty acid-derived metabolites that synergize with antibiotics by reducing fatty acid uptake. We identified two independent Tn insertions in *sigB* that increased drug tolerance. SigB is an alternative sigma factor that was reported to be required for mycobacterial tolerance to RIF and INH (75, 76). Our results contrast with these studies, possibly due to our use of different growth media, and suggest that SigB can under certain conditions limit *M. tuberculosis* drug tolerance.

Our results highlight several advantages of screening low-complexity Tn mutant pools made from an arrayed library. These include identification of Tn mutants with robust phenotypes from selection conditions with strict bottlenecks, efficient recovery of individual Tn mutants, and reproducibility of mutant phenotypes upon individual retesting. However, our results also uncovered one drawback of this method: the potential for recovery of Tn mutants with secondary mutations that alter the phenotype of interest. In standard Tn-seq screens that use high-complexity Tn mutant libraries, secondary mutations are less likely to influence identification of genes that significantly impact fitness due to the presence of multiple independent Tn insertions in each gene. Our screen identified numerous Tn mutants with decreased drug tolerance, but it is possible that many of these strains harbor spontaneous secondary mutations causing loss of PDIM production, similar to the *clpS*::Tn and *ppgK*::Tn mutants. Distinguishing whether the decreased drug tolerance of these mutants is due to the Tn insertion or loss of PDIM will require recovery and individual retesting of these Tn mutants, which can be efficiently done from our arrayed Tn mutant library.

Overall, our results demonstrate that *M. tuberculosis* requires PDIM for drug tolerance under nutrient starvation conditions *in vitro*. As PDIM is also a critical virulence determinant that is required to counteract host immune pressures, our results suggest that inhibitors of PDIM production could synergize with both host-imposed stress and existing antibiotics to kill *M. tuberculosis* more efficiently. This could dramatically shorten TB treatment times and prevent emergence of new drug-resistant strains. PDIM biosynthesis is a complex process, requiring multiple polyketide synthases, fatty acyl ligases and thioesterases, several of which have already been explored as potential drug targets (62, 77, 78). It will be critical to determine whether PDIM deficiency also increases *M. tuberculosis* antibiotic susceptibility during infection to further support development of new inhibitors targeting PDIM production, which we intend to pursue in our future studies.

## METHODS

### Bacterial strains and culture conditions

Bacterial strains used in this study are listed in Table S5. For routine culture, *M. tuberculosis* Erdman wild-type and derivative strains were grown aerobically at 37°C in Middlebrook 7H9 (Difco) liquid medium supplemented with 10% albumin-dextrose-saline (ADS), 0.5% glycerol, and 0.1% Tween-80 or on Middlebrook 7H10 (Difco) agar supplemented with 10% oleic acid-albumin-dextrose-catalase (OADC; BD Biosciences) and 0.5% glycerol. Frozen stocks of *M. tuberculosis* strains were made from mid-exponential phase cultures by adding glycerol to 15% final concentration and storing at −80°C. All experiments used MtbYM rich liquid medium (MtbYM), pH 6.6 (38) supplemented with 10% OADC and 0.05% tyloxapol. For P_i_ starvation experiments, bacterial were grown in P_i_-free MtbYM (made by replacing Na_2_HPO_4_ and KH_2_PO_4_ with NaCl and KCl and buffering with 50 mM 3-(*N*-morpholino)propanesulfonic acid (MOPS), pH 6.6). Antibiotics were used at the following concentrations unless otherwise noted: kanamycin (Kan) 25 μg/ml for agar or 15 μg/ml for liquid, hygromycin (Hyg) 50 μg/ml, ciprofloxacin (CIP) 8 μg/ml, rifampicin (RIF) 0.1 μg/ml, and isoniazid (INH) 0.2 μg/ml.

### Creation and mapping of a *M. tuberculosis* Erdman arrayed transposon mutant library

Transposon (Tn) mutagenesis of wild-type *M. tuberculosis* Erdman was performed by transduction with the mycobacteriophage phAE159 carrying the *Himar1* Tn as previously described (79). Wild-type bacteria were grown to mid-exponential phase (OD_600_ of 0.4-0.6) in 7H9 broth, washed and resuspended in MP buffer (50 mM Tris pH 7.6, 150 mM NaCl, 10 mM MgCl_2_, 2mM CaCl_2_), then transduced with mycobacteriophage at a MOI of at least 20:1 for 3 hours at 40°C. Phage adsorption was stopped with Stop Buffer (MP buffer with 60 mM Na-citrate and 0.6% Tween-80) and transduced cells were plated on MtbYM agar pH 6.6 with Kan at a density of 100-200 colonies per plate. Plates were incubated at 37°C with 5% CO_2_ for at least 3 weeks. Approximately 8000 individual Tn mutant colonies were picked from plates into 600 μl of MtbYM broth in 1 ml v-bottom Matrix screw-cap tubes in a 96-well rack (Thermo Scientific) and incubated with shaking at 37°C for 2 weeks, until turbid.

Tn mutants were orthogonally pooled using the Straight Three strategy and sequenced, as previously described (39). Briefly, the Tn library was pooled in 2 groups of 40 racks each (racks 1-40 and 41-80). For each rack, a small volume of culture was removed from each tube and combined appropriately to form 8 row pools (rows A-H), 12 column pools (columns 1-12) and a rack pool. Each individual rack pool was aliquoted into one sample for sequencing and nine 1 ml aliquots for experimental use, which were stored at −80°C with glycerol at 15% final concentration. After pooling, glycerol was added at 15% final concentration to each Tn mutant culture and racks were stored at −80°C. For each group of 40 racks, analogous row and column pools were pooled from all 40 plates to generate 8 row and 12 column samples. These were multiplexed with the 40 individual rack pools for a total of 60 samples for Tn-seq sequencing. Genomic DNA (gDNA) was extracted from each row, column and rack pool using the CTAB-lysozyme method (80) and submitted to the University of Minnesota Genomics Center (UMGC) for library creation and Tn-seq sequencing as described below. Tn mutants associated with the reads were traced back to their rack location in the arrayed Tn library using two approaches: Straight Three (39) and Knockout Sudoku (40).

### Drug tolerance Tn-seq screen in stationary phase and P_i_-starvation

Five frozen rack pools (1 ml each) generated during orthogonal pooling were inoculated in 250 ml MtbYM broth with Kan and grown at 37°C with aeration to mid-exponential phase (OD_600_ of 0.4-0.6). A portion of the culture was removed to start a 250 ml P_i_-starved culture. Bacteria were washed twice in P_i_-free MtbYM broth, inoculated in P_i_-free MtbYM with Kan at OD_600_ = 0.1 and incubated at 37°C with aeration for 72 hours. The remaining MtbYM culture was grown at 37°C with aeration for a total of 7 days to reach early stationary phase. We experimentally determined that at least 10^6^ CFU of WT Erdman are recovered from a 12 ml culture after 9 days of drug treatment in either P_i_-free or stationary phase conditions (Fig 1). Therefore, as input controls, the P_i_-free or stationary phase cultures were serially diluted and plated on MtbYM agar at a density of ~10^6^ CFU/plate before addition of antibiotics. Cultures were then split into triplicate 12 ml antibiotic treated (CIP+INH or RIF+INH) or untreated-control cultures and incubated with aeration at 37°C for 9 days. Antibiotic-treated bacteria were collected by centrifugation (3720 x *g*, 10 min), washed twice with an equal volume of PBS-T (Gibco PBS, pH 7.4 with 0.05% Tween-80) to remove antibiotics, concentrated 100-fold in PBS-T, and plated on YM agar with Kan to recover at least 10^6^ CFU. Untreated control cultures were serially diluted and plated at a density of ~10^6^ CFU/plate. Plates were incubated at 37°C with 5% CO_2_ until the biomass on the agar was confluent, up to two weeks. Confluent plates were flooded with 2 ml of GTE buffer (80) and gently scraped with a plastic 10 μl loop to loosen the biomass. Bacteria were collected by centrifugation (3720 x *g*, 10 min) and gDNA was extracted from cell pellets by the CTAB-lysozyme method (80) and cleaned using the Genomic DNA clean and concentrator kit (Zymo) before submitting to UMGC for Tn-seq library preparation and Illumina sequencing.

### Transposon sequencing (Tn-seq) and data analysis

Tn-seq was performed as previously described (38). *M. tuberculosis* genomic DNA was fragmented with a Covaris S220 ultrasonicator and a whole genome library was prepared using the TruSeq Nano library preparation kit (Illumina). Library fragments containing Tn junctions were PCR-amplified from the whole genome library using the Tn-specific primer Mariner_1R_TnSeq_noMm and Illumina p7 primer (Table S6). The amplified products were uniquely indexed to allow sample pooling and multiplexed sequencing. Resulting Tn-seq libraries were sequenced on an Illumina 2500 High-output instrument in 125-bp paired-end output mode using v4 chemistry (Illumina). Sequencing reads were filtered to remove reads without the Tn sequence “GGACTTATCAGCCAACCTGT”. The 5’ Illumina adaptor sequences were trimmed using BBDuk (https://sourceforge.net/projects/bbmap/). Each trimmed read was cut to 30 bases and sequences not starting with TA were removed. Remaining reads were mapped to the *M. tuberculosis* Erdman genome (NC_020559.1) using HISAT2. Mapped reads were counted at each TA insertion site in the *M. tuberculosis* Erdman genome to generate read count tables for TnseqDiff analysis. TnseqDiff normalized the read counts using the default trimmed mean of M values (TMM) normalization method (81, 82) and then determined conditional essentiality for each TA insertion site between experimental conditions (control/input, CIP+INH/input, RIF+INH/input, CIP+INH/control, RIF+INH/control). TnseqDiff calculated the fold change and corresponding two-sided *P*-value for each TA insertion site (41). All *P*-values were adjusted for multiple testing using the Benjamini-Hochberg procedure in TnseqDiff. The cut-off values for statistical significance were set at a fold-change of > log_2_ ±2 and an adjusted *P*-value < 0.025.

### Recovery of Tn mutants from the arrayed Tn mutant library

Each Tn mutant individually retested was isolated from the tube corresponding to the Tn mutant location in the arrayed library by streaking for individual colonies on MtbYM agar containing Kan. Plates were incubated for at least 3 weeks at 37°C. Up to four individual colonies were picked and grown in 10 ml of MtbYM broth with Kan at 37°C with aeration until turbid. The Tn insertion site was confirmed by PCR using a gene-specific primer 5’ or 3’ of the TA site and a primer specific to the Tn Kan-resistance cassette (Table S6) followed by Sanger sequencing.

### Tn mutant complementation

To complement Tn mutants, we generated an integrating vector encoding Hyg resistance, pMV306hyg, by replacing the Kan-resistance cassette in pMV306 (83) with a Hyg-resistance cassette. pMV306 lacking the Kan-resistance cassette was PCR-amplified with primers pMV306_F and pMV306_R containing SbfI and AflII restriction sites, respectively, and digested with SbfI and AflII. A Hyg-resistance cassette was removed from pJT6a (84) by restriction with SblI and AflII, gel purified and ligated to the the SbfI and AflII-digested pMV306 to create pMV306hyg (Table S5).

Complementation vectors were constructed by PCR-amplifying each gene along with ~300 bases 5’ of the translation start site to include the native promoter (Table S6). PCR products were cloned in pCR2.1 TOPO (Invitrogen) and sequenced, then removed from pCR2.1 by restriction digestion with XbaI and HindIII and ligated to XbaI and HindIII-digested pMV306hyg. The pMV361hyg-*ppsD* complementation vector was generated by replacing the Kan-resistance cassette pMV361-*ppsD* (63) with a Hyg-resistance cassette by Gibson assembly. Linearized pMV361-*ppsD* without the Kan-resistance cassette and the Hyg-resistance cassette from pMV306hyg were generated by PCR with primer pairs pMV361.FOR/pMV361.REV and hyg_fwd_2/hyg_rev_2, respectively (Table S6), and gel purified. PCR products were assembled with NEB Hifi assembly master mix (New England Biolabs) following the standard protocol, and Hyg-resistant transformants were selected. The *ppsD* gene and Hyg-resistance cassette in pMV361hyg-*ppsD* were Sanger sequenced. Cosmid 8C3.1 containing the genomic region *rv1317-rv1343* surrounding *clpS* was obtained from the lab of William R. Jacobs. Tn mutants were electroporated with the corresponding complementation vector or cosmid as described (85). Transformants were selected on Middlebrook 7H10 agar containing Kan and Hyg. The presence of the complementing plasmid or cosmid was confirmed by PCR (Table S6).

### Individual retesting of Tn mutant antibiotic tolerance

Bacteria were grown from frozen stocks to mid-exponential phase (OD_600_ of 0.4-0.7) in 7H9 complete medium. For P_i_-free experiments, starter cultures were washed twice with P_i_-free MtbYM broth, resuspended at an OD_600_ of 0.1 in P_i_-free MtbYM broth and grown for 72 hours. For stationary phase experiments, starter cultures were diluted to an OD_600_ of 0.05 in MtbYM broth and grown for 7 days. For exponential phase experiments, starter cultures were diluted to an OD_600_ of 0.025 in MtbYM medium, incubated for 3 days to reach mid-exponential phase (OD_600_ of 0.4-0.7), then diluted to OD_600_ of 0.2. Cultures were serially diluted and plated on 7H10 agar to enumerate the input CFU/ml. Cultures were then split into CIF+INH-treated or RIF+INH-treated and aliquoted to create 12 ml single-use cultures for each time point. This method enables formation of stable drug tolerant populations by reducing production of toxic reactive oxygen species that are generated due to changes in oxygen saturation upon repeated sampling of a culture (15). Cultures were washed with PBS-T, serially diluted and plated on 7H10 agar to determine surviving CFU/ml as previously described (86). Plates were incubated at 37°C with 5% CO_2_ for at least 4 weeks prior to enumerating surviving CFU.

### Minimum inhibitory concentration (MIC) assay

Bacteria were grown from frozen stocks to mid-exponential phase (OD_600_ ~0.5) in MtbYM broth and diluted to OD_600_=0.01 in 5 ml fresh MtbYM broth. Antibiotics were added at 2-fold increasing concentrations. Cultures without antibiotics were included as controls. Cultures were incubated at 37°C with aeration. The OD_600_ of each culture was measured on day 7 for INH or day 14 for RIF and CIP. The MIC_90_ was defined as the minimum concentration of antibiotic required to inhibit at least 90% of growth relative to the no antibiotic control.

### Whole-genome sequencing of *M. tuberculosis* wild-type Erdman and Tn mutants

Genomic DNA (gDNA) was extracted from the wild-type Erdman parental strain and *rv0457c*::Tn, *ppgK*::Tn and *clpS*::Tn mutants grown to late-exponential phase in 7H9 broth by the CTAB-lysozyme method (80). The gDNA was cleaned with the Genomic DNA concentrator and cleanup kit 25 (Zymo) then sheared to ~300 bp by ultrasonication (Covaris S220). Sizing post-shearing was done with an Agilent Bioanalyzer. Libraries were prepared from sheared gDNA using the NEBNext Ultra II DNA Library Prep Kit for Illumina (New England Biolabs). Briefly, the ends of the fragmented DNA were repaired by 5’ phosphorylation and dA-Tailing, followed by Illumina adaptor ligation. Adaptor-ligated DNA was size-selected for a 350 bp insert using AMPure beads (Beckman Coulter). Adaptor-ligated DNA was then PCR-amplified, uniquely barcoded, and cleaned with AMPure beads. Library QC, pooling and Illumina sequencing was done at the UMGC. Samples were sequenced on an Illumina iSeq 100 with 150 bp paired-end output. To generate a consensus sequence for each strain, paired reads were mapped to the *M. tuberculosis* Erdman reference genome (NC_020559.1) using the “map to reference” function in Geneious 2020 software (Biomatters, Ltd.) with the following settings: mapper – Geneious; sensitivity – medium sensitivity/fast; fine tuning – iterate up to 5 times.

To identify single nucleotide polymorphisms (SNPs) in Tn mutant genomes, the WT Erdman and Tn mutant consensus sequences were aligned in Geneious using the “Align Whole Genomes” function with the default Mauve Genome parameters (Alignment algorithm – progressiveMauve algorithm; automatically calculate seed weight; compute Locally Colinear Blocks (LCBs); automatically calculate minimum LCB score; full alignment). Regions containing PE-PGRS genes that were poorly mapped in either consensus sequence were excluded from SNP analysis. SNPs identified in genes required for PDIM synthesis were confirmed by PCR amplification and Sanger sequencing (Table S6). To confirm Tn insertion sites in the Tn mutants, reads were mapped to the *Himar1* Tn sequence in Geneious as described above. Sequences adjacent to the Tn were compared to the *M. tuberculosis* Erdman reference genome to identify the Tn insertion site.

### Phthiocerol dimycocerosate (PDIM) labeling and detection

To detect PDIM production, we used a radiolabeling and thin layer chromatography (TLC) method (63). Briefly, *Mtb* cultures grown to mid-logarithmic phase in 10 ml of 7H9 broth were labeled for 48 hr with 10 μCi of [1-^14^C] propionic acid, sodium salt (American Radiolabeled Chemicals, Inc; specific activity 50-60 mCi/mmol). Labeled bacteria were collected by centrifugation (2500 x *g*, 10 min). Apolar lipids were extracted twice in 2 ml 10:1 (vol:vol) methanol:0.3% NaCl and 2 ml petroleum ether by vortexing for 4 min, collecting the upper petroleum ether layer after phase separation by centrifugation (750 x *g*, 10 min). Combined petroleum ether fractions were inactivated for 1 hour with an equal volume of chloroform, then evaporated overnight to reduce the extract to ~4 ml. Extracts (30 μl) were spotted on a silica gel 60 F_254_ TLC plate (5×10 cm; Supelco). The TLC plate was developed in 9:1 (vol:vol) petroleum ether:diethyl ether, air dried and exposed to a phosphor storage screen (Amersham) for 3 days. Radioactive bands were detected using a Typhoon FLA 9500 Imager (GE Healthcare). Intensity of the radioactive signal in each PDIM spot was quantified with ImageJ.

### Custom scripts and code

All customs scripts and R code used for Illumina sequence read processing and TnseqDiff data analysis are available at https://github.com/bloc0078/umn-tischler-tnseq.

### Statistical analysis

A student’s unpaired t-test (two-tailed) was used for pairwise comparisons between WT, mutant and complemented strains. *P* values were calculated using GraphPad Prism 8.0 (GraphPad Software, Inc). *P* values < 0.05 were considered significant.

### Data availability

Raw sequencing data from Tn-seq and whole genome sequencing will be made publicly available in FASTA format through the Data Repository for the University of Minnesota.

## ACKNOWLEDGEMENTS

This work was completed in part with resources, including instrumentation and staff, at the University of Minnesota Genomics Center (UMGC, RRID: SCR_012413) and the University of Minnesota University Imaging Centers (UIC, RRID: SCR_020997). We thank Drs. Yusuke Minato and Anthony Baughn for developing the custom MtbYM rich media and for helpful discussions. Cosmid 8C3.1 was provided by Dr. William R. Jacobs, Jr. This work was supported by a National Institutes of Health Director’s New Innovator Award DP2AI112245 from NIAID (ADT), T90 DE02273 from NIDCR (AM Block) and a University of Minnesota Doctoral Dissertation Fellowship (SNB). The content is solely the responsibility of the authors and does not reflect the official views of the National Institute of Dental & Craniofacial Research, the National Institutes of Allergy and Infectious Disease or the National Institutes of Health.

## SUPPLEMENTAL LEGENDS

**Fig S1. Genome distribution and mapping confidence of the *M. tuberculosis* Erdman arrayed Tn library.** A) Heat map showing distribution of distances between Tn insertions sites for all unambiguously mapped Tn mutants. B) Number of Tn mutants unambiguously mapped per tube in library racks 1-40 (black) or 41-80 (gray).

**Fig S2. Mutants with altered fitness upon drug treatment in P_i_-limited MtbYM medium compared to the no drug control.** (**A-B**) Volcano plots of TnseqDiff statistical analysis of Tn-seq data for P_i_ starved Tn mutant pools treated with CIP+INH (**A**) or RIF+INH (**B**) compared to the no drug control. Dashed lines indicate ± 2 Log_2_ fold change and adjusted *p*-value < 0.025 statistical significance cutoffs. Tn mutants meeting significance are colored. (**C-D**) Venn diagrams displaying the number of Tn mutants with significant negative (**C**) or positive (**D**) fold changes in relative fitness.

**Fig S3. Mutants with altered fitness upon drug treatment during stationary phase in MtbYM medium compared to no drug control.** (**A-B**) Volcano plots of TnseqDiff statistical analysis of Tn-seq data for Tn mutant pools grown to stationary phase in MtbYM and treated with CIP+INH (**A**), or RIF+INH (**B**) compared to the no drug control. Dashed lines indicate ± 2 Log_2_ fold change and adjusted *p*-value < 0.025 statistical significance cutoffs. Tn mutants meeting significance are colored. (**C-D**) Venn diagrams displaying the number of Tn mutants with significant negative (**C**) or positive (**D**) fold changes in relative fitness.

**Fig S4. Whole-genome sequencing results for *rv0457c*::Tn, *ppgK*::Tn, and *clpS*::Tn mutants.** (**A-C**) Sample of whole-genome sequencing reads from the contigs for *rv0457c*::Tn (**A**), *ppgK*::Tn (**B**), and *clpS*::Tn (**C**) confirming *Himar1* Tn insertion at the predicted TA site. Black arrows indicate the TA site predicted by Tn-seq library mapping. Black boxes surround portions of the sequencing read containing the *Himar1* Tn sequence. (**D-E**) Whole-genome sequencing reads aligned to show single nucleotide polymorphisms in *ppsE* for *ppgK*::Tn (**D**) and *ppsD* for *clpS*::Tn (**E**) compared to WT.

**Table S1.** Mapped locations of Tn mutants in the arrayed *Mycobacterium tuberculosis* Erdman Tn library.

**Table S2.** Raw read counts for each TA Tn insertion site for all P_i_ starvation and stationary phase drug tolerance Tn-seq screens.

**Table S3.** Complete TnseqDiff statistical analysis of Tn-seq results for all P_i_ starvation and stationary phase drug tolerance Tn-seq screens.

**Table S4.** Tn mutants with statistically significant fold changes in relative abundance determined by TnseqDiff analysis for all P_i_ starvation and stationary phase drug tolerance Tn-seq screens.

**Table S5.** Plasmids, cosmid, and strains used in this study.

**Table S6.** Oligonucleotide primers used in this study.

