## Supplemental Figures for "*Mycobacterium tuberculosis* requires the outer membrane lipid phthiocerol dimycocerosate for starvation-induced antibiotic tolerance"

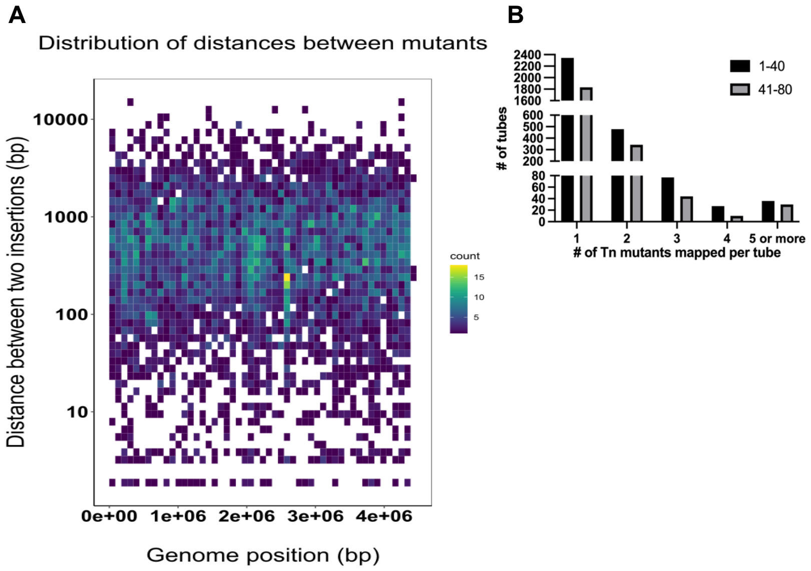


**Fig S1. Genome distribution and mapping confidence of the *M. tuberculosis* Erdman arrayed Tn library.** A) Heat map showing distribution of distances between Tn insertions sites for all unambiguously mapped Tn mutants. B) Number of Tn mutants unambiguously mapped per tube in library racks 1-40 (black) or 41-80 (gray).

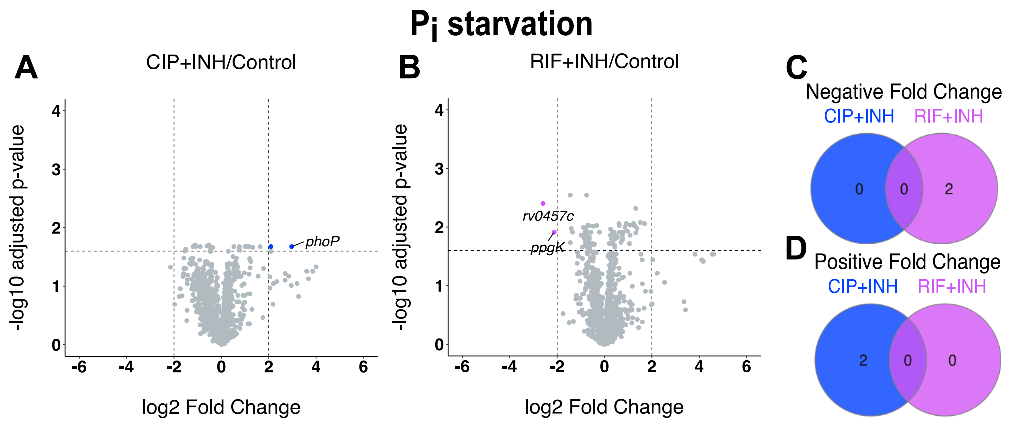


**Fig S2. Mutants with altered fitness upon drug treatment in $P_i$-limited MtbYM medium compared to the no drug control.** (A-B) Volcano plots of TnseqDiff statistical analysis of Tn-seq data for $P_i$ starved Tn mutant pools treated with CIP+INH (**A**) or RIF+INH (**B**) compared to the no drug control. Dashed lines indicate $\pm$ 2 $Log_2$ fold change and adjusted *p*-value < 0.025 statistical significance cutoffs. Tn mutants meeting significance are colored. (**C-D**) Venn diagrams displaying the number of Tn mutants with significant negative (**C**) or positive (**D**) fold changes in relative fitness.

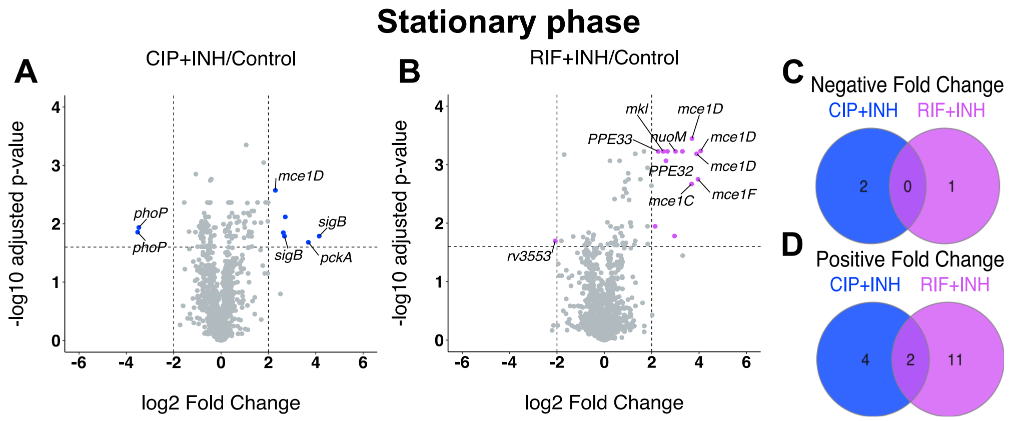


**Fig S3. Mutants with altered fitness upon drug treatment during stationary phase in MtbYM medium compared to no drug control.** (**A-B**) Volcano plots of TnseqDiff statistical analysis of Tn-seq data for Tn mutant pools grown to stationary phase in MtbYM and treated with CIP+INH (**A**), or RIF+INH (**B**) compared to the no drug control. Dashed lines indicate ± 2 $Log_2$ fold change and adjusted *p*-value < 0.025 statistical significance cutoffs. Tn mutants meeting significance are colored. (**C-D**) Venn diagrams displaying the number of Tn mutants with significant negative (**C**) or positive (**D**) fold changes in relative fitness.

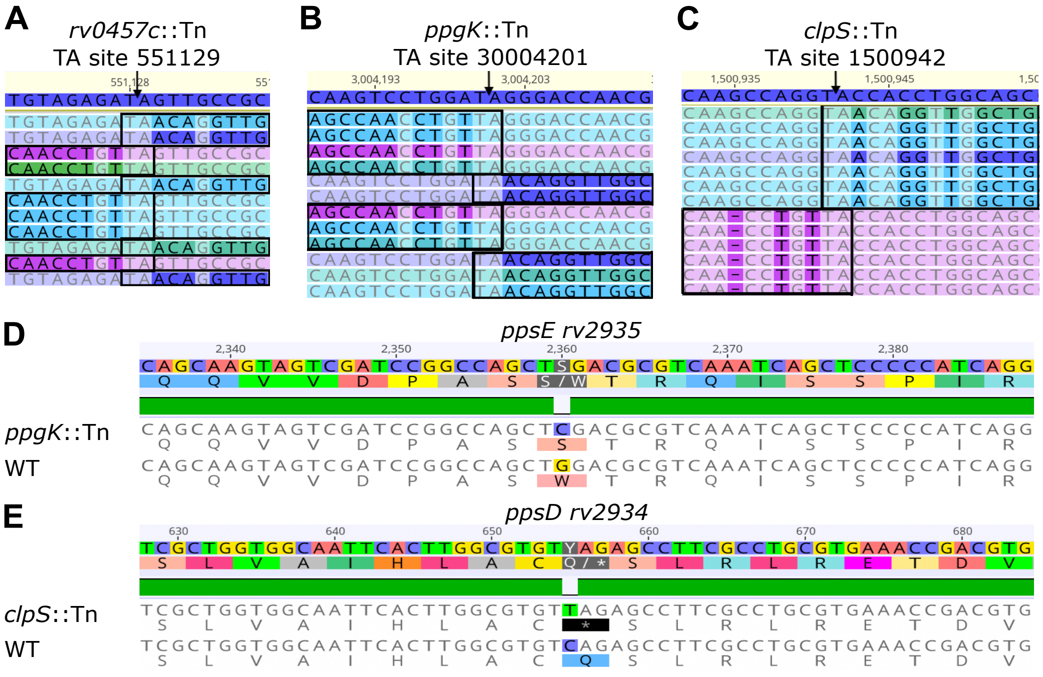


**Fig S4. Whole-genome sequencing results for *rv0457c*::Tn, *ppgK*::Tn, and *clpS*::Tn mutants.** (**A-C**) Sample of whole-genome sequencing reads from the contigs for *rv0457c*::Tn (**A**), *ppgK*::Tn (**B**), and *clpS*::Tn (**C**) confirming Himar1 Tn insertion at the predicted TA site. Black arrows indicate the TA site predicted by Tn-seq library mapping. Black boxes surround portions of the sequencing read containing the Himar1 Tn sequence. (**D-E**) Whole-genome sequencing reads aligned to show single nucleotide polymorphisms in *ppsE* for *ppgK*::Tn (**D**) and *ppsD* for *clpS*::Tn (**E**) compared to WT.
